# Extrinsic Noise Suppression in Micro RNA mediated Incoherent Feedforward Loops

**DOI:** 10.1101/422394

**Authors:** Alberto Carignano, Sumit Mukherjee, Abhyudai Singh, Georg Seelig

## Abstract

MicroRNA mediated incoherent feed forward loops (IFFLs) are recurrent network motifs in mammalian cells and have been a topic of study for their noise rejection and buffering properties. Previous work showed that IFFLs can adapt to varying promoter activity and are less prone to noise than similar circuits without the feed forward loop. Furthermore, it has been shown that microRNAs are better at rejecting extrinsic noise than intrinsic noise. This work studies the biological mechanisms that lead to extrinsic noise rejection for microRNA mediated feed forward network motifs. Specifically, we compare the effects of microRNA-induced mRNA degradation and translational inhibition on extrinsic noise rejection, and identify the parameter regimes where noise is most efficiently rejected. In the case of static extrinsic noise, we find that translational inhibition can expand the regime of extrinsic noise rejection. We then analyze rejection of dynamic extrinsic noise in the case of a single-gene feed forward loop (sgFFL), a special case of the IFFL motif where the microRNA and target mRNA are co-expressed. For this special case, we demonstrate that depending on the time-scale of fluctuations in the extrinsic variable compared to the mRNA and microRNA decay rates, the feed forward loop can both buffer or amplify fluctuations in gene product copy numbers.

## I. Introduction

Stochasticity in gene expression is known to be modulated by biological feedback and feed-forward network motifs [1]. In particular, recent work has found that IFFLs - a network architecture where an upstream regulator directly activates a downstream target and also indirectly inhibits it - are capable of buffering against changes in promoter activity, as well as reducing the stochasticity in gene expression [2]-[4]. Furthermore, it has also been shown that the reduction of noise is more effective at the post-transcriptional level for IFFLs [5]. In this study we focus on understanding the noise rejection properties of a particular class of post-transcriptionally regulated IFFLs, where the negative regulatory link is implemented with a microRNA (miRNA).

MiRNA are a class of non-coding, regulatory RNAs that have been linked with the post-transcriptional regulation of important biological processes including differentiation, development and disease [6]. As was the case for transcriptionally regulated IFFLs, experimental studies with miRNA mediated IFFLs have shown that they have effective noise-suppressing and buffering properties in some parameter regimes, often displaying interesting nonlinear behaviors [2], [4]. miRNA regulation of gene expression is is believed to occur through two different regulatory modes: i) by degrading mRNAs that the miRNAs bind to, and ii) by preventing efficient translation of miRNA bound mRNAs into proteins (translation-inhibition) [7]. To elucidate the mechanisms underlying these properties, several mathematical models have been proposed to describe this class of IFFLs [8]-[10]. However, these models either rely on large mathematical nonlinearities [8], or on complex network architecture [10] to explain the nonlinear noise-rejection properties of the system. Such models also often make modeling assumptions that are not justified using biological mechanisms. On the other hand, data-fitted models [9] don’t explore the full parameter space, and risk to overlook important components in the machinery. Hence, a qualitative, parameter-independent model that could provide an easy biological explanation is currently missing. Here, we propose a simple quasi-linear model that utilizes mechanisms inferred from previous experimental studies [4], [7] and mild assumptions on the underlying extrinsic noise sources. Through these basic assumptions, we explain the previously-reported biological behaviors, and recapitulate the mechanisms at the origin of the noise-repression mechanism with particular attention to their biological interpretation. These insights allow us to precisely pinpoint the qualitative contribution of each cellular machinery on the overall behavior, providing a framework for understanding current biological systems and for designing synthetic systems for noise rejection.

We initially focus on the the more biologically common situation of static extrinsic noise, which arises when the mRNA degradation occur on a much faster time scale than than that set by the cell cycle. We analytically obtain conditions for extrinsic noise rejection for general miRNA mediated IFFLs, demonstrating that the noise correlation alone defines the parameter space for noise rejection. Then, we demonstrate that high mRNA-miRNA binding rate could lead to an overall noise increase, and give a simple biological explanation. Moreover, we show that, contrary to what was previously reported [9], translation-inhibition is a key-player in reducing protein noise. Finally, we extend the analysis to dynamic extrinsic noise, focusing on the special case where the mRNA and miRNA are derived from the same gene (single-gene IFFL). We demonstrate that even for this simple system, dynamic extrinsic noise can lead to a wide range of noise rejection regimes depending on the relative stability of the mRNA and the miRNA.

## II. Model of IFFL System

We now outline the miRNA-mRNA IFFL model that we use in this study, starting from the underlying biochemical reactions. mRNA (*M*) and miRNA (S) are transcribed from genes g_1_ and g_2_ respectively at constant rates *α*_*m*_, *α*_*s*_. The process of translation then converts mRNA into protein (P) at constant rate *α*_*p*_; miRNA, by definition, is not coding for any protein. To complete the reaction network, we add the degradation processes:

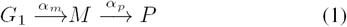

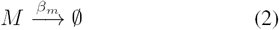

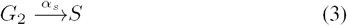

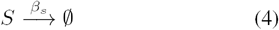

The miRNA-based regulation proceeds primarily through two groups of steps namely miRNA mediated degradation and translational inhibition which affect each of these reactions separately. Many models for miRNA based degradation have been proposed [7], [11]-[13]. For the purposes of this work we rely on a previously-adopted model [4]: we assume that mRNA and miRNA form an irreversible complex C that is then degraded into the miRNA alone at rate k_c_. Moreover, we assume that mRNA can be translated into protein even when bound to the miRNA, although at a lower rate. This factor accounts for the miRNA-induced translation inhibition that has been previously reported [7], [9]:

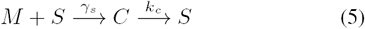

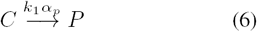

where *k*_1_ is a real number between 0 and 1, where *k*_1_ = 0 implies full translation inhibition (no protein from the complex) and *k*_1_ = 1 implies that the complex has the same translation rate as the mRNA alone.

We now write a mathematical model describing 1 and 5: the species concentrations (or copy numbers) are represented with the corresponding lower case bold-faced variables which represent random processes.

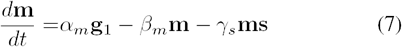

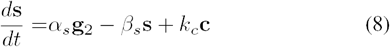

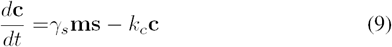

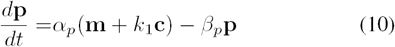

The production rates of mRNA and miRNA are dependent respectively on the positive random variables g_1_ and g_2_, which correspond to the average number of active genes. Genes g_1_ and g_2_ can be transcribed either dependently (*Cov*(g_1_, g_2_) ≠ 0) or independently (*Cov*(g_1_, g_2_) = 0).

This model can be further simplified by assuming that the mRNA-miRNA complex formation reaches steady state at a faster pace than the other processes. Hence, we get 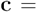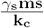. This simplifies equations 8-10 to:

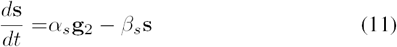

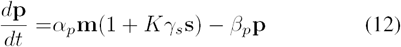

Where 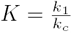. In the special case where there is complete translational inhibition we observe that *K* = 0.

Steady state mRNA and protein levels of the open loop system 1 can easily be computed:

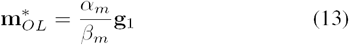

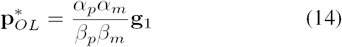

Under the reasonable assumption that there is negligible variability in the cellular processes within a cell population, the extrinsic noise source is the variability of the transcription of the genes g_1_ and g_2_. This enables us to avoid making any assumptions on the distributions of the parameter values. The coefficient of variations of the open loop system are given by:

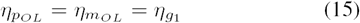

The corresponding steady state levels in the general IFFL system according to 11 are:

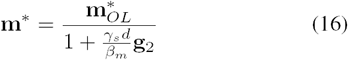

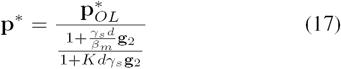

Where 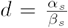. Equations 16 and 17 highlight the separate dependency of the system equilibria on the open-loop steady states and the miRNA regulation.

## III. Static Extrinsic Noise Rejection in General IFFL Systems

### Extrinsic Noise Measurement at Steady State for Static Noise

Extrinsic noise arises from diversity in cell populations, such as differences in cell size, in uptake of a external signal, or in the cell cycle phase. The two random variables g_1_ and g_2_ representing gene expression level, account for the extrinsic noise in our model. We consider two cases: when the noise is static in nature (g_1_ and g_2_ are fixed distributions over time) and when it is time-varying.

To quantify steady-state noise rejection in microRNA-based systems, we use the square of the coefficient of variation defined as the ratio:

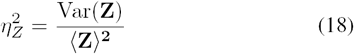

where Z is the random variable of interest, and Var ( ) and ⟨•⟩ are the standard notation for the variance and expected value operators. The coefficient of variation allows to compare noise in processes that have different means. This is essential in our study, as microRNA-control reduces the level of mRNA and protein.

We use a first-order Taylor-Delta approximations to obtain the ratio and products of the two random variables (gi and g_2_), which leads to the following expression for the coefficient of variation 18:

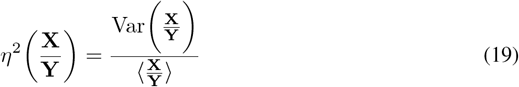

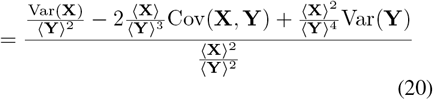

This expression can be simplified to:

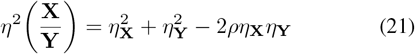

where *ρ* is the Pearson product-moment correlation coefficient between the two random variables **X** and **Y**.

We can now compute the coefficient of variations for the mRNA (16) and protein (17) level as:

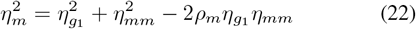

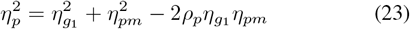

where we define the miRNA-dependent noise contribution for mRNA (*η*_*mm*_) and protein (*η*_*pm*_) as:

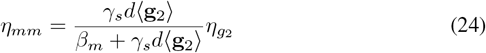

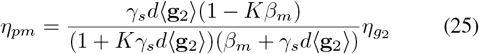

and *ρ*_*m*_ and *ρ*_*p*_ as:

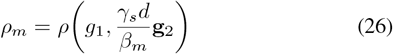

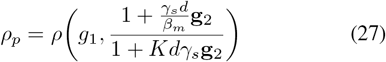

According to formulae 22 and 23, miRNA regulation introduces noise in the system (represented by the quantities 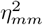 and 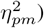), but it also cancels part of the noise components of g_1_ if the two processes are positively correlated (*ρ >* 0). The noise cancellation terms, 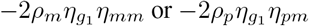 suggest that the noise is canceled from both sources depending on how well they correlate with each other.

### Conditions for noise cancellation in miRNA-regulated IFFLs

To understand the extrinsic noise reduction properties of miRNA regulation, we studied the values of *η*_*mm*_ and *η*_*pm*_ in 22 and 23 that satisfy condition: 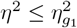, when the noise of the close-loop system is lower than for the open loop. It is easy to see that a necessary condition for noise rejection is *ρ*_*m*_ > 0 and *ρ*_*p*_ > 0. At the mRNA level (eq 24), this condition is guaranteed iff *ρ*(g_1_, g_2_) > 0, which occurs when the same upstream gene positively regulates both g_1_ and g_2_, as is the case in an IFFL. Conversely, uncorrelated miRNA or coherent feed forward loops (CFFLs) do not have extrinsic noise cancellation properties at the mRNA level according to our model.

At the protein level (eq 25), two conditions need to be satisfied to have *ρ*_*p*_ > 0:

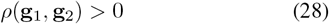

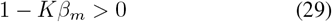

Condition 28 is the same as for mRNA, which implies that only IFFLs can achieve noise rejection at the protein level. Condition 29 guarantees that the miRNA-mRNA complex does not translate into protein faster than mRNA alone would be degraded. If this were to happen, more protein would be produced from the complex than from the open loop mRNA. Therefore the miRNA regulation would not repress protein expression, but instead facilitate it. Hence, translation-inhibition (*K* > 0) could cancel the noise-rejection property of the IFFL.

Even if ρ > 0, the parabolic form of 22 and 23 implies that there is a limited range of values of *η*_*mm*_ and *η*_*pm*_ that leads to extrinsic noise rejection, before the extra noise introduced by the miRNA machinery leads to worse performance than in the open loop (Figure1). The limiting case is when 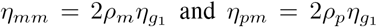, where the amount of noise that is introduced by the miRNA is equivalent to the amount of canceled noise (*η* = *η*_g1_). The minimum of the parabola corresponds to the maximum noise rejection that could be achieved. This is reached for 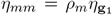 and 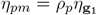, for which the noise rejection ratio is:

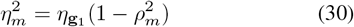

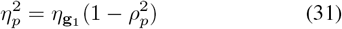

Thus, optimal noise rejection is achieved for perfect correlation (*ρ* = 1).

In summary, we showed that miRNA-based IFFLs are the only network architecture that can lead to noise reduction in simple miRNA mediated systems. We also demonstrated that correlation between the two types of noise (miRNA-based and mRNA) is the main factor in extrinsic noise cancellation. However, very high miRNA-related noise (ή_*mm*_ > 2*p*_*m*_ ή_*g1*_ or ^®^_*pm*_ > 2*p*_*p*_ ^®^_*g*1_ increases the system noise even at maximum correlation (*ρ*_*m*_ = 1 and *ρ*_*p*_ = 1), as one would expect (Figure 1).

**Fig. 1.**
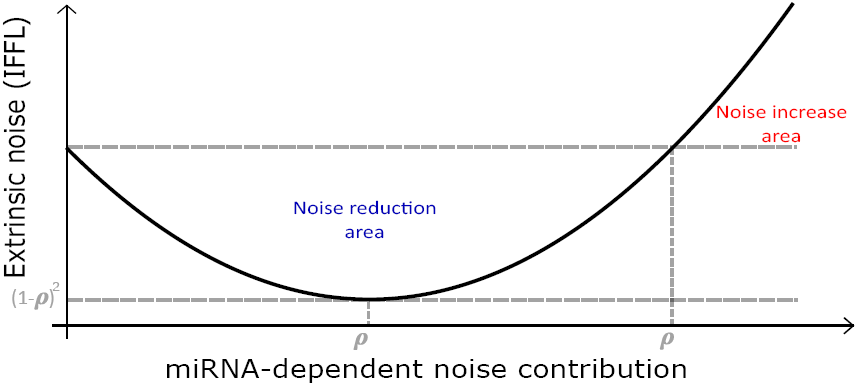
Extrinsic noise rejection in microRNA based IFFLs.

### High miRNA-mRNA binding rate could lead to increased extrinsic noise

We investigated the relationship between the miRNA-mRNA binding rate (γ_*s*_) and the extrinsic noise rejection properties of the IFFLs. We showed in the previous subsection that the amount of noise introduced by the miRNA regulation 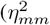 or 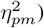 could be balanced by noise cancellation (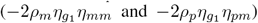. Hence, we computed the value of *η*_*mm*_ at the mRNA and the protein level and studied their dependencies to γ_*s*_. It could be shown that *η*_*mm*_ in 24 has a monotonic dependency on γ_*s*_:

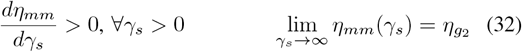

This equations shows that there is a maximum amount of noise that could be introduced by the miRNA regulation to the system at the mRNA level, and this is bounded by the amount of noise in the input signal g_2_ (specifically, by 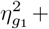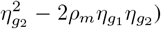). The mRNA extrinsic noise tends to this value in the limit, although for high values of γ_*s*_, **m\*** → 0. In this range, the mRNA dynamics are entirely defined by fluctuations in the miRNA, as all mRNA are in a complex (Figure2(a)). Moreover, if *n*_*g*2_ < 2*p*_*m*_*n*_*g*1_, according to 22, noise rejection is achieved for all values of γ_*s*_.

The optimal noise rejection is reached for:

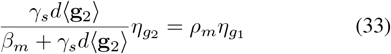

This equation shows that the mRNA noise is reduced when the miRNA-induced degradation can compete with the natural mRNA degradation *β*_*m*_;. Increasing or reducing the value of *β*_*m*_ requires a higher or lower value of γ_*s*_ to reach the minimum of the parabola. At the protein level, translation inhibition introduces a non-trivial nonlinear dynamic. In fact, *η*_*pm*_ is not monotonic on γ_*s*_, and it reaches a maximum for:

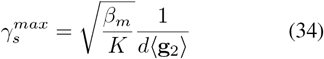

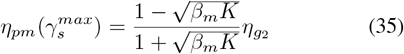

We notice that the maximum value of 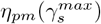 is positive iff 1 - *β*_*m*_ *K* > 0, which is assured by the noise-rejection condition 29. Equation 25 also shows that the value of *η*_*pm*_ as γ_*s*_ → ∞ is 0. Hence the maximum amount of noise that could be introduced by the miRNA regulation to the system at the protein level is reached for 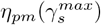 (Figure 2(b)). This quantity is less or equal than 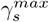, satisfying the equality for *K* = 0 (complete translation-inhibition), when the dynamic of the protein noise is equivalent to the mRNA one.

Depending on the values of 35, there are three possible scenarios:

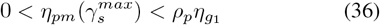

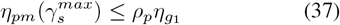

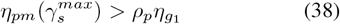

**Fig. 2.**
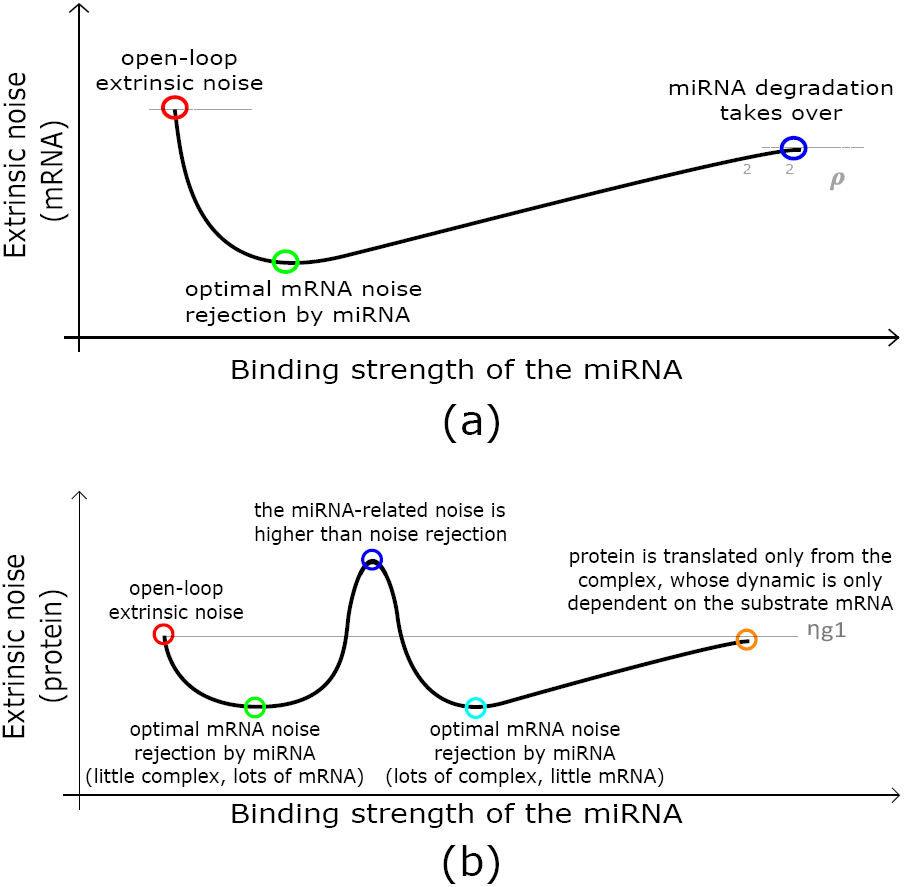
High miRNA-mRNA binding rate has a nonlinear effect on the overall extrinsic noise. (a)Noise rejection at the mRNA level for a miRNA-based IFFL. miRNA-mRNA binding strength drives noise reduction from 0 (red circle) to a minimum (green circle, maximum noise cancellation). Higher binding rates introduce more miRNA noise in the mRNA spectrum till it reaches its maximum (blue circle).(b) Noise rejection at the protein level for a miRNA-based IFFL. Protein is generated from both complex and mRNA (assuming *K* ≠ 0): for intermediate values of γ_*s*_, this leads to a biphasic distribution, which increases the overall noise (dark blue circle).

If 36 is satisfied, then there is only one optimal binding rate γ_*s*_, which is sub-optimal for the given distribution g_1_ and g_2_. If 37 is satisfied, then optimal noise rejection is reached for a unique value of γ_*s*_. If 38 is satisfied, then optimal noise rejection is reached for two different values of γ_*s*_, and noise increases between the two (local maximum). Moreover, if is bigger than 2*p*_*p*_*n*_*g*1_, then this local maximum is higher than the open loop noise. In all three scenarios, the IFFL noise tends to the open-loop noise as γ_*s*_ → ∞ (Figure2(b).

In summary, the protein noise follows the mRNA noise closely for low binding rates. As the binding rate increases, the miRNA-mRNA complex becomes dominant and competes with mRNA for translation to protein: these two separate translation events lead to a bimodal protein population (’threshold effect’), which increases the total extrinsic noise (maximum highlighted in blue in Figure 2(b)). At higher binding rate, the miRNA-mRNA complex mostly dominates translation: protein noise is then dependent on the miRNA noise, reaching a second minimum (highlighted in light blue in Figure 2(b)). For even higher values of γ_*s*_, protein translation occurs mainly through the miRNA-mRNA complex, because little free mRNA is left. In this regime, the protein noise is independent on miRNA, since at steady state the complex is dependent only on the amount of the substrate mRNA. Hence, *η*_*p*_ reaches the open-loop mRNA noise *r*_*s*_ → ∞ 1. as γ_*s*_ → ∞.

### Translation-inhibition modulates the functional range of miRNA extrinsic noise rejection

To address the importance of translation-inhibition for the miRNA-induced noise rejection, we analyzed the dependency of our results in the previous section to perturbation of the translation-inhibition parameter *K.* Equations 34 and III show that for a fixed value of γ_*s*_, the performance of the miRNA-IFFL could be tuned by increasing or decreasing the translation-inhibition parameter *K* or the mRNA degradation *β*_*m*_. However, tuning these parameters have opposite effects on optimality: 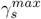 is reduced if *K* is increased, and it increases if *β*_*m*_ is increased. This is not surprising since the two mechanisms are competing for mRNA degradation.

As shown in Figure 3, increasing *K* changes the location and height of the noise peak both in intensity (y-axis) and in its corresponding binding strength (x-axis). On the other hand, if *K* = 0, the dynamic follows the mRNA noise closely and reaches its maximum at 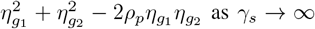 as γ_*s*_ → ∞ (green line in Figure 3).

**Fig. 3.**
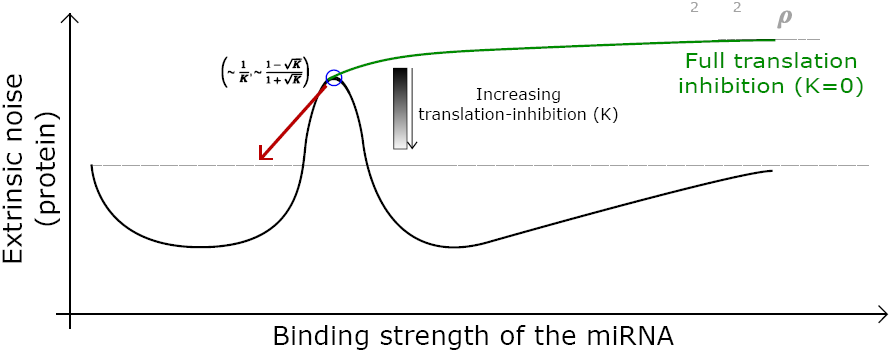
Translation-inhibition modulates the functional range of miRNA extrinsic noise rejection. Upon binding to mRNA, miRNA interferes with the protein translation machinery (translation-inhibition, *K* ≠ 0). This effect modulates the extrinsic noise peak (blue circle) both in intensity (y-axis) and in its corresponding binding strength (x-axis). For high values of *K* (low translation-inhibition), overall noise rejection occurs at a wider range of γ_*s*_ values rather than for full translation inhibition (*K* = 0).

Hence, translation-inhibition modulates the noise-rejection range of the miRNA-mRNA binding rate γ_*s*_. For high-noise miRNA, full translation-inhibition (*K =* 0) reduces protein noise over a larger range of γ_*s*_ (since 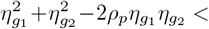), while little translation-inhibition is preferred for low-noise miRNA.

## IV. Static and Dynamic Extrinsic Noise Rejection in sgFFL Systems

### A. Static noise rejection by sgFFLs

A very common and functionally important sub-class of IFFLs is the single gene feed forward loop or sgFFL [14], [15]. sgFFLs are a special class of IFFLs where the mRNA and the miRNA are co-expressed by the same gene [4], [16]. Typically, the miRNA is located within an intron of the gene encoding the mRNA. The mRNA and the miRNA are first transcribed together into one transcript and then separated from each other by splicing [17], [18]. Mathematically, this can be expressed as g_1_ = g_2_ = g, which simplifies expression 22 to

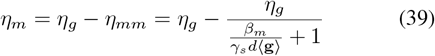

As a result of this simplification, the mRNA noise *η*_*m*_ can be reduced arbitrarily close to 0 by increasing 7_S_, since *η*_*mm*_ → *η*_*g*_ as γ_*s*_ → ∞. Extrinsic variations of mRNA levels would be canceled out by equivalent variations of miRNA: each cell would then be reduced to the same mRNA steady state, defined by the fixed biochemical rates of degradation. This behavior is conserved at the protein level if there is full translation inhibition. If one assumes that some of the complex gets translated, then there is an optimal value of γ_*s*_ for which extrinsic noise is minimized, and the protein concentration is not reduced to 0. This value is equivalent to:

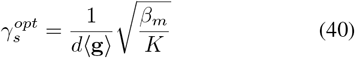

which is the sgFFL case of 34. This values is a ratio between mRNA that escapes miRNA degradation (naturally degraded at rate *β*_*m*_ or translated as miRNA-mRNA complex) and total amount of miRNA (d⟨ g⟩).

### B. Dynamic noise rejection by sgFFLs

Having so far studied the effect of static extrinsic noise on sgFFL circuits, we now consider the scenario of dynamic noise by allowing

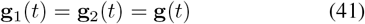

to be a stochastic process modeled as an Ornstein-Uhlenbeck (OU) process

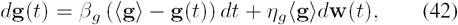

where ⟨ g ⟩ is the steady-state mean level of g(*t*), w(*t*) is a Wiener process, *η*_*g*_ and *β*_*g*_ are the magnitude (coefficient of variation) and time-scale of fluctuations in g(*t*), respectively. This stochastic process drives the sgFFL circuit described earlier by the following system of different equations

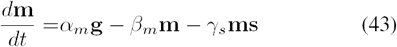

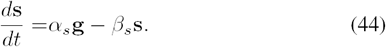

For simplicity, in this subsection we will only focus on noise at the mRNA level, and hence ignore the protein dynamics. Note that the limit *β*_*g*_ → 0, where fluctuations in g(*t*) are significantly slower than the mRNA-miRNA dynamics, recovers the scenario of static noise that we have up till now considered. A key question of interest is as we now vary *β*_*g*_ what miRNA-mRNA interaction strengths provide the most efficient noise buffering?

To address these questions, we investigate the statistical moments of the joint stochastic processes {**g**(*t*), **m**(*t*), **s**(*t*)}. Readers are referred to [19], [20] for details on deriving moment dynamics for hybrid systems of the form (42)-(44) that couple stochastic and ordinary differential equations. It turns out that the nonlinear product term **ms** in (43) makes computations non-trivial due to unclosed moment dynamics - the time evolution of the lower-order moments always depends on high-order moments. Moving forward, one general approach is to exploit closure schemes that approximate higher-order moments as nonlinear functions of lower-order moments [21]-[35]. Here we use an alternative approach based on linearizing the nonlinearity

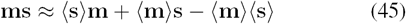

where ⟨m⟩ and ⟨s⟩ denote the mean steady-state levels for the mRNA and miRNA, respectively. Using this approximation in place of ms in (43) result in a linear dynamical system driven by an OU process, and standard theory can now be applied to compute moments. More specifically, let *µ* denote a vector consisting of all the first and second order moments of {g(*t*), m(*t*), s(*t*)}, then its time evolution is given by the linear system

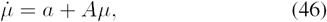

where vector *a* and matrix *A* depend on model parameters [19], [20]. Steady-state analysis of (46) yields a complicate formula for the steady-state noise (coefficient of variation) in mRNA levels *η*_*m*_, as given by (47) on top of the next page. As expected, in the limit of static noise (*β*_g_ → 0), (47) reduces to (39), where 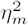 monotonically decreases with increasing mRNA-miRNA interaction strength *γ*_*s*_.

In the limit of no mRNA-miRNA interaction, the mRNA noise level is given by

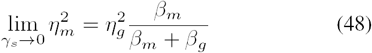

and is an increasing function of the mRNA decay rate *β*_*m*_. Intuitively, as we decrease the mRNA half-life, it becomes less efficient in averaging out upstream fluctuations in g(*t*), and more noise from g(*t*) propagates to the mRNA level. In the limit of strong mRNA-miRNA interaction,

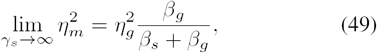

and comparing (50) and (48) reveals an intriguing result: for sufficiently fast fluctuation in g(*t*), such that, 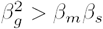

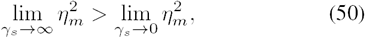

i.e., strong mRNA-miRNA feed forward interaction will actually amplify noise compared to the open loop system. This counter intuitive result can be understood from the fact that while the incoherent feed forward loop cancels external noise, it also decreases the mRNA half-life which causes more noise propagation from g to m. The balance of these two effects results in an interesting set of behaviors illustrated in Fig 4 which plots (47) as a function of γ_*s*_. In particular, we see

**Fig. 4.**
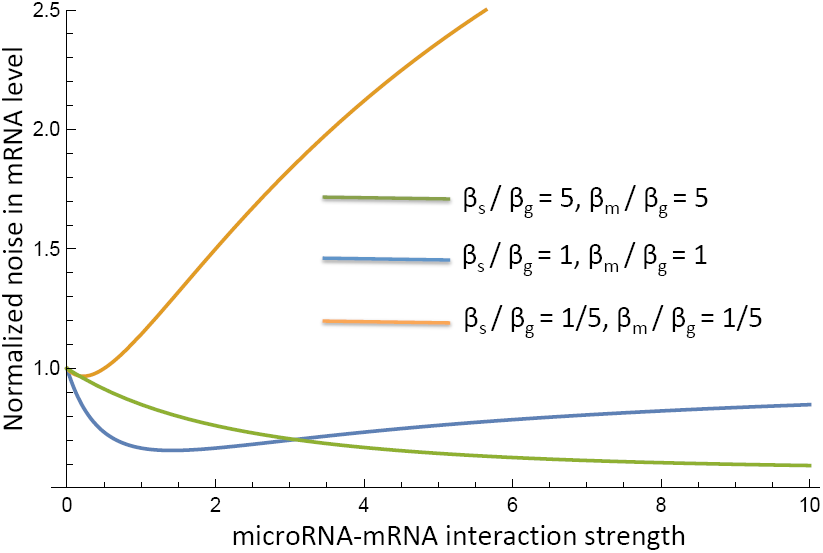
Dynamical noise rejection by miRNA mediated IFFLs. Plot of noise in mRNA levels as given by (47) as a function of the mRNA-miRNA interaction strength for different values *β*_*g*_ (time-scale of extrinsic fluctuations), *β*_*m*_ (mRNA decay rate) and *β*_*s*_ (miRNA decay rate). While low values of *β*_*g*_ cause *η*_*m*_ to monotonically decrease with γ_*s*_., (green line), values of *β*_*g*_ comparable or higher than *β*_*m*_ and *β*_*s*_ result in a nonmonotonic profile for *η*_*m*_ (blue and orange lines), where mRNA noise is minimized at an optimal interaction strength. Note that in this plot mRNA noise levels are normalized by their value in the open loop system with no mRNA-miRNA interaction.

- For slow fluctuations in g(*t*), mRNA noise level decreases with increasing Y_S_ as expected form our analysis of static noise (green line).
- When time-scale of fluctuations in g(*t*) are comparable to mRNA and miRNA half-lives, 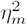 first decrease with increasing γ_*s*_ to reach a minimum, and then increase with γ_*s*_ to create a U-shape profile (blue line).
- For sufficient fast fluctuations in g(*t*), the U-shape become shallower and shifts to the left such that 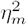 is mostly an increasing function of γ_*s*_ (orange line).

In summary, our results show that if the time-scale of fluctuations in the external variable is somewhat comparable to the system dynamics, then introducing noise statically in the system can lead to erroneous results. Given the space constraints, we have only focused on the noise at the mRNA level in sgFFLs, and future work investigating dynamic noise propagation at the protein level in general mRNA-miRNA feed forward circuits is clearly warranted.

### V. Conclusion and Future Work

Here we have identified the regimes for noise rejection at the mRNA and the protein level for miRNA-mediated IFFLs. In the case of static extrinsic noise, we investigated how the different modes of post-transcriptional regulation by miRNAs affect noise levels. Interestingly, we found that IFFLs are the only microRNA mediated process with extrinsic noise reduction capability. Furthermore, we found that due to translational inhibition, the regimes for noise rejection are quite different at the level of mRNA and protein. While ordinarily this added non-linearity leads to an increase in noise, under certain biologically realistic circumstances, this effect can be reversed leading to a greater range of noise reduction at the protein level. We also mathematically verify the experimental observation [4] that high mRNA-miRNA binding rate can lead to an increase in extrinsic noise.

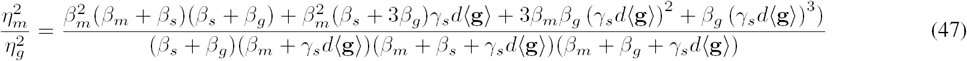

For the case of dynamic extrinsic noise, we limit our investigation to the special case of co-expressed mRNA and miRNA (sgFFL system). Our results show counter-intuitive effects if the time-scale of fluctuations in g(t) are comparable or faster than the time scale of mRNA and miRNA turnover. In particular, while for static extrinsic noise the mRNA noise is predicted to monotonically decrease with increasing mRNA-miRNA interaction strength, in the case of dynamic noise (for all other model parameters fixed) there exists an optimal feed forward strength that minimizes mRNA noise.

Future work will be aimed at extending this analysis to intrinsic noise. Such a study will enable a more comprehensive understanding of the noise rejection properties of such systems. Another promising direction of future work relates to studying the noise modulation by more complicated miRNA mediated network motifs, particularly those involving multiple correlated or uncorrelated miRNAs.

